# Ayurvedic *Amalaki Rasayana* promotes improved stress tolerance and thus has anti-aging effects in *Drosophila melanogaster*

**DOI:** 10.1101/050476

**Authors:** Vibha Dwivedi, Subhash C. Lakhotia

## Abstract

**Ethnopharmacological relevance:** *Amalaki Rasayana* (AR) is a common Ayurvedic herbal formulation of *Phyllanthus emblica* fruits and other ingredients and is used for general good health and healthy aging. We earlier reported it to improve life history traits and to suppress neurodegeneration as well as induced apoptosis in *Drosophila*.

**Aim of the study:** To examine effects of dietary AR supplement on cell stress responses in *Drosophila melanogaster*.

**Materials and methods:** Larvae/flies, reared on normal food or on that supplemented with 0.5% (w/v) AR, were exposed to crowding, thermal or oxidative stress and examined for survival, stress tolerance and levels of lipid peroxides, SOD and HSPs.

**Results:** Wild type larvae/flies reared on AR supplemented food survived the various cell stresses much better than those reared on normal food. AR-fed mutant *park*^*13*^ or *DJ-1β*^*Delta93*^ (Parkinson’s disease model) larvae, however, showed only partial or no protection, respectively, against paraquat-induced oxidative stress, indicating essentiality of *DJ-1β* for AR mediated oxidative stress tolerance. AR feeding reduced the accumulation of reactive oxygen species (ROS) and lipid peroxidation even in aged (35 day old) wild type flies while enhancing superoxide dismutase (SOD) activity. We show for the first time that while Hsp70 or Hsp83 expression under normal or stress conditions was not differentially affected by AR feeding, Hsp27 levels were elevated in AR fed wild type control as well as heat shocked larvae. Therefore, besides the known anti-oxidant activity of *Phyllanthus emblica* fruits, dietary AR also enhances cellular levels of Hsp27.

**Conclusion:** In the context of the reported “anti-aging” and “healthy-aging” effects of AR, the present in vivo study on a model organisms shows that AR feeding significantly improves tolerance to a variety of cell stresses through reduced ROS and lipid peroxidation and enhanced SOD activity and Hsp27. Such improved cellular defences following dietary AR provide better homeostasis and thereby improve the life-span and quality of organism’s life.

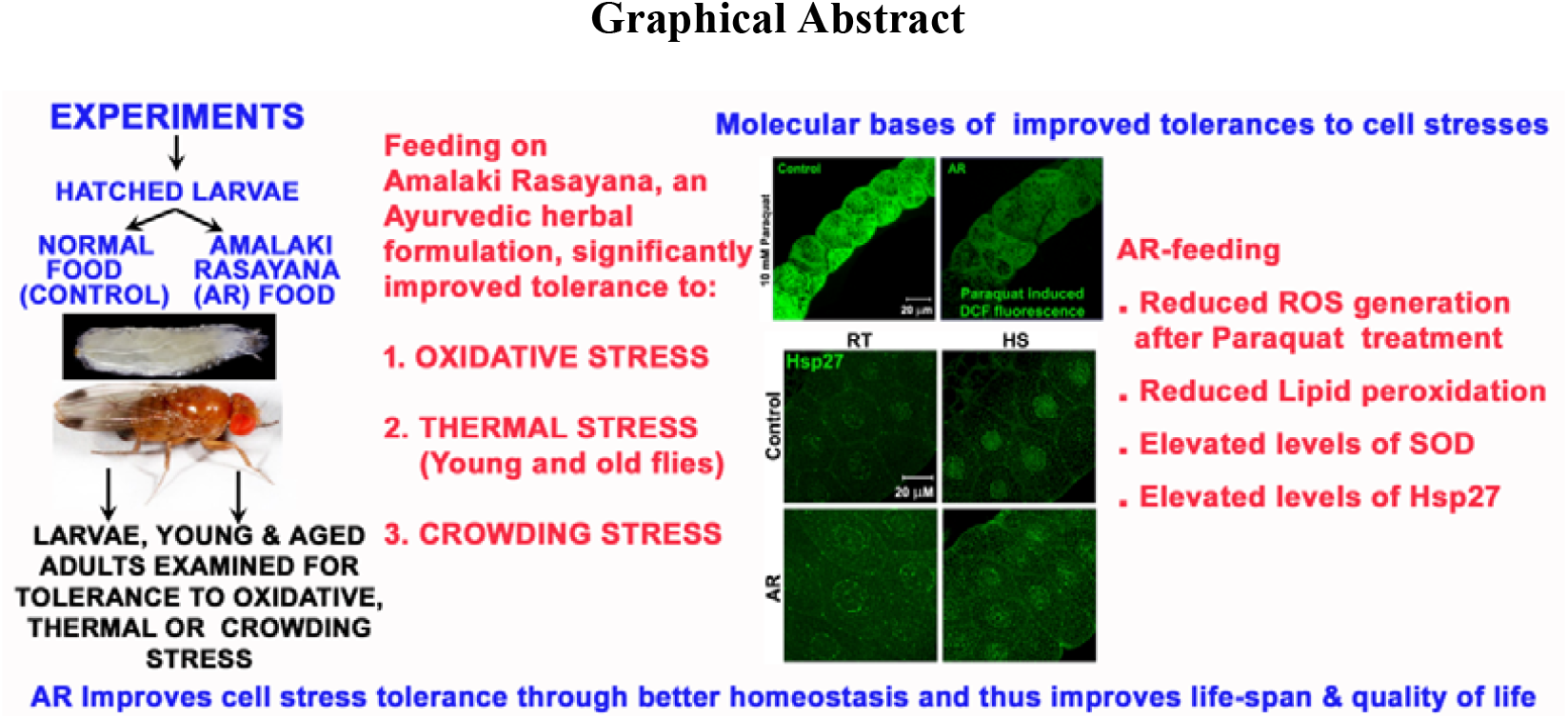

## Introduction

The ancient Ayurvedic literature as available today does not elaborate bases and mechanisms of actions of Ayurveda’s varied therapeutic preparations and procedures in terms of our contemporary understanding of biology and other sciences. Therefore, a new class of studies was initiated to rigorously apply the methods of modern science to understand its major concepts, procedures and mechanistic aspects (Valiathan, 2006). To this end, we are using the fruit fly, *Drosophila melanogaster*, as a model organism (Dwivedi et al. 2012, 2013, 2015) to get insight into the cell biological and biochemical bases of actions of *Amalaki Rasayana* (AR), which is one of the prominent herbal formulations described in Ayurvedic classics like *Charak Samhita* (Sharma, 1994) and *Ashtang Hridaya* (Murthy, 2000). AR continues to be widely used in view of the claim that it enhances life expectancy, body strength, intellect, fertility and reduces age-related debilities (Puri, 2003). *Amalaki Rasayana* is prepared from fruits of Amla or Indian gooseberry (*Phyllanthus emblica*, synonym *Emblica officinalis*) with some other ingredients through a specific and elaborate process (see Dwivedi et al. 2012). We reported earlier (Dwivedi et al. 2012) that, in general agreement with AR’s suggested therapeutic applications in Ayurveda (Puri, 2003), rearing of wild type *Drosophila melanogaster* on food supplemented with 0.5% AR improved various biological parameters like life-span, fecundity, stress-tolerance etc. We further reported that dietary AR suppressed neurodegeneration in fly models of polyQ-disorders or Alzheimer’s disease (AD) without any side-effects (Dwivedi et al. 2013) and that it substantially inhibited induced but not developmental apoptosis (Dwivedi et al. 2015).

In order to effectively meet the continuous challenges by the dynamic external and internal environmental conditions to the homeostasis of cells and the organism, biological systems have evolved a variety of highly conserved cell stress responses which attempt to restore homeostasis or induce cell death to minimize damage to the organism (Arya et al. 2007; Ekengren et al. 2001; Evgen’ev et al. 2014; Feder and Hoffmann, 1999). One of the factors that bring about aging and age-related debility is the progressively declining ability to effectively respond to the variety of cell stresses experienced by organism as part of its normal life (Arya et al. 2007; Haigis and Yanker, 2015; Phillips et al. 1989; Wang et al. 2013; Yan, 2014; Zhang et al. 2015). Since AR is believed to provide for healthy aging, in this study we examined if rearing of wild type flies on AR supplemented food affects their tolerance to applied stressors like crowding, severe hyperthermia and increased reactive oxygen species (ROS) production. Growth under crowded conditions increases metabolic waste levels which have detrimental effects on survival of the organism (Borash and Ho, 2001; Chippindale et al. 1993; Mueller and Barter, 2015). Thermal stress affects many important cellular processes like DNA replication, transcription, post-transcriptional processing, transport and translation in diverse organisms (Arya et al. 2007; Feder and Hoffmann, 1999; Lindquist, 1986). Wild type laboratory populations of *Drosophila melanogaster* larvae or flies exposed to temperatures >37^0^C, often get immobilized or “knocked down” and show a high incidence of delayed lethality, which increases with age due to a less efficient stress response (Calderwood et al. 2009; Landis et al. 2012; Morrow and Tanguay, 2003). The normally generated pro-oxidant reactive oxygen species (ROS) or free radicals significantly increase upon exposure to adverse physicochemical, environmental or pathological conditions. The resulting oxidative stresses adversely affect cellular activities through DNA damage, modifications of polypeptides, lipids etc (Adelman et al. 1988; Lushchak, 2014; Sies, 2015; Zhao and Haddad, 2011). A variety of pathological conditions, including cancers, autoimmune diseases, and neurodegenerative diseases also involve generation of ROS and mitochondrial dysfunction (Gandhi and Abramov, 2012; Greene et al. 2005; Melkani et al. 2013; Shukla et al. 2014). Oxidative damage is considered to be one of the major factors underlying aging process (Landis et al. 2012; Schieber and Chandel, 2014; Wang et al. 2013; Yan, 2014; Zhang et al. 2015). Our results show that dietary AR significantly improves tolerance of young as well as aged wild type flies to crowding, hyperthermic and paraquat-induced oxidative stress. However, *park*^*13*^ and *DJ-1β*^*Delta93*^ mutant flies (Parkinson’s disease model) showed only partial or no improvement, respectively, in oxidative stress tolerance following AR feeding. The AR-fed wild type aged flies showed reduced levels of hydroperoxides and lipid-peroxidation while levels of SOD and Hsp27 were elevated. It appears, therefore, that dietary AR improves fly’s tolerance to diverse stresses through multiple paths, the improved stress tolerance in turn promotes increase in their longevity and fecundity as noted earlier (Dwivedi et al. 2012). In view of the fact that the fly model is increasingly used in recent years for understanding human health issues (Perrimon et al. 2016), the present findings provide a mechanistic basis for the use of AR in traditional Ayurvedic practices for promoting healthy life and healthy aging.

## Materials and Methods

### 1.1 *Drosophila* stocks and rearing

The *Oregon R*^+^ strain of *Drosophila melanogaster* was used as wild type. The following two mutant stocks, *w*; +/+*; park*^*13*^/*Tm6B*, and *w*^*1118*^; +/+*; DJ-1β*^*Delta93*^ stocks (Bloomington stock no 33601), associated with Parkinson’s disease model in flies, were also used. The *park*^*13*^ is a null allele of *parkin* in *Drosophila* (Greene et al. 2005) while *DJ-1 β*^*Delta93*^ is a null allele of *DJ-1β* (Meulener et al. 2005). The *Hsc70Cb-YFP* stock (no. 115570 from DGGR, Kyoto) expressing YFP-tagged Hsc70Cb was also used.

All fly stocks were reared on standard agar-cornmeal-sugar-yeast *Drosophila* food at 24^0^C±1^0^C. *Amalaki Rasayana* was prepared by Arya Vaidya Sala (Kottakkal, Kerala, India) following a well-defined and quality-controlled process which has been described earlier (Dwivedi et al. 2012). It was mixed with the standard food (0.5% w/v) for rearing of experimental larvae and/or flies at 24^0^C±1^0^C as described earlier (Dwivedi et al. 2012), with parallel controls reared on the regular un-supplemented food. In all experiments, eggs were collected from fly stocks that had always been reared on the regular food. For each experiment, the regular (control) and the formulation (AR) supplemented foods were prepared from the same batch and likewise all larvae/adults for a given experiment were derived from a common pool of eggs and reared in parallel on the regular or AR supplemented food.

### 1.2 Crowding resistance assay

Freshly hatched *Oregon R*^+^ larvae were reared in standard food vials, each containing about 5g of 0.5% AR supplemented or regular (control) food. 25 (normal density), 50 (moderately crowded) or 100 (highly crowded) freshly emerged larvae were transferred to each food vial and allowed to grow and eclose. The median life span of flies was calculated (Dwivedi et al. 2012) in each case.

### 1.3 Thermal Tolerance Assay

Flies of the desired age (3, 15, 30, 45 or 60 day after eclosion), reared at 24±1^0^C either on regular or on 0.5% AR supplemented food through the larval stages were lightly etherised and transferred to plastic vials (25 flies per vial). The flies were allowed to recover from anaesthesia for at least 4 h in food vials before subjecting them to heat shock (HS) in empty vials in a water bath maintained at 38°C for 60 min. The number of flies that could not fly or move and fell down (knocked down) in the vial during the period of HS was monitored every 15 min (Dwivedi et al, 2012). After 60 min at 38^0^C, the flies were transferred to food vials at 24±1^0^C and the numbers of flies surviving after 24 h were recorded. Eight replicates of 25 flies each were examined for each experimental condition.

### 1.4 Paraquat induced oxidative stress

Male flies of the *Oregon R*^+^ (wild type), *w*; +/+*; park*^*13*^/*Tm6B*, or *w*^*1118*^; +/+*; DJ-1β*^*Delta93*^ stocks of desired age, reared either on formulation supplemented or regular food since first instar larval stage, were starved in empty vials for 6 hours before treatment with N, N′-dimethyl −4, 4′-bipyridinium dichloride or paraquat (Sigma-Aldrich, India). Flies were transferred to a food-free vial containing a filter paper soaked in 10mM paraquat in 5% sucrose solution and kept in a moist chamber (Hao et al. 2007). The flies were monitored for survival at 12 h intervals till all the flies in a vial died. Parallel control flies were kept in vials with filter paper soaked in 5% sucrose solution only. Eight replicates of 25 flies each were examined for each experiment.

### 1.5 Reactive Oxygen Species (ROS) estimation

To assay cellular ROS levels, the desired tissues of wild type late third instar larvae, 1 or 35 day old wild type flies fed on regular or AR supplemented food since the first instar stage were dissected out in 1X PBS and incubated in 1µg/ml 2’,7’-dichlorfluorescein-diacetate (DCFH-DA) at 24^0^C (DCFD, Sigma-Aldrich, India) for 5 min followed by washing twice in 1X PBS and immediate viewing in the LSM 510 Meta Zeiss confocal microscope. Conversion of the non-fluorescent DCFH-DA to highly fluorescent compound 2’,7’-dichlorfluorescein (DCF) is directly proportional to levels of hydroperoxides in cells (Cathcart et al. 1983). Therefore, the total fluorescent intensity of DCF indicates the level of ROS in the sample.

### 1.6 Lipid peroxidation measurement

35 day old 10 male and 10 female wild type flies that were fed on formulation supplemented food or on regular food since the first instar larval stage were homogenized in 150 µl 1X phosphate buffered saline (PBS, 13mM NaCl, 0.7mM Na_2_HPO_4_, 0.3mM NaH_2_PO_4_, pH 7.0) at 4^0^C. Lipid peroxidation was measured following the method of Ohkawa et al. (1979) and levels of lipid peroxides were expressed in terms of nmoles of malondialdehyde (MDA) formed/hour/mg of protein.

### 1.7 Superoxide dismutase Assay

35 day old 10 males and 10 wild type females, fed on formulation supplemented or regular food since first instar stage, were homogenized in 150 µl extraction buffer (100 mM potassium phosphate and 100 mM EDTA, pH 7.5) at 4^0^C. The Cu–Zn SOD activity was estimated following Kakkar et al. (1984) and expressed as specific activity (enzyme units/min/mg protein)

### 1.8 Immunoblotting and immunostaining of tissues

Western blots of total larval proteins from larvae fed on regular food or AR supplemented food, and subjected to desired experimental condition, were challenged with rat anti-Hsp70 (7Fb, 1:1000 dilution, Velazquez and Lindquist, 1984) or rat anti-Hsc/Hsp70 (7.10.3, 1:500 dilution, Palter et al. 1986) or anti-β-tubulin (E7, 1:200 dilution, Sigma Aldrich, India). The primary antibody binding was detected using alkaline phosphatase conjugated anti-rat or anti-mouse IgG (Bangalore Genei, India) secondary antibody, respectively as described earlier (Prasanth et al. 2000). For immunostaining, the desired tissues were dissected out in Poels’ salt solution (PSS) (Lakhotia and Tapadia, 1998) and transferred to freshly prepared 3.7% paraformaldehyde for 20 min at RT and processed further for immunostaining as described (Prasanth et al. 2000). The different primary antibodies used were: (1) rat anti-Hsp70 (7Fb, 1:100 dilution), (2) mouse monoclonal anti-Hsp27 (ab49919, 1:100 dilution, Abcam, UK), (3) mouse monoclonal anti-Hsp90 (SPA 830, 1:100 dilution, Stressgen, USA). Appropriate secondary antibodies conjugated either with Cy3 (1:200, Sigma-Aldrich) or Alexa Fluor 488 (1:200, Molecular Probes) were used to detect the given primary antibody. The immunostained tissues were counterstained with DAPI, mounted in DABCO and examined under LSM510 Meta Zeiss laser scanning confocal microscope using appropriate laser, dichroic and barrier filters. Quantitative analysis and colocalization of immunofluorescence were carried out using the Histo and Profile tools in the LSM510 Meta software.

All images were assembled using Adobe Photoshop 7.0.

### 1.9 RNA isolation and Real Time-PCR

Third instar larvae reared either on regular food or on AR supplemented food were dissected out after heat shock/recovery in PSS and total RNA was isolated using the TRI Reagent as per the manufacturer’s (Sigma-Aldrich, India) instructions. RNA pellets were resuspended in nuclease free water. The cDNA was synthesized and real-time quantitative PCR (RT-qPCR) was carried out as described previously (Singh and Lakhotia, 2016), using appropriate primers and SYBR-Green dye on 7500 Real Time PCR System (Applied Biosystems) with the 7500 software v2.0.4. The primers used were: (i) G3PDH: Forward: 5′-CCACTGCCGAGGAGGTCAACTA-3′, Reverse: 5′-GCTCAGGGTGATTGCGTATGCA-3′, (ii) Hsp70: Forward: 5′-AGGGTCAGATCCACGACATC-3′, Reverse: 5′-CGTCTGGGTTGATGGATAGG-3′, (iii) Hsp27: Forward: 5′-GTCCATGCCCACGATCTGTT-3’, Reverse: 5′-CGACACATCCATGCACACCT-3’, (iv) Hsp83 Forward: 5′-CCTGGACAAGATCCGCTATG-3’, Reverse: 5′-GAAACCCACACCGAACTGAC-3’.

### 1.10 Statistical Analysis

Sigma Plot 11.0 was used for statistical analyses. All percentage data were subjected to arcsine square root transformation and expressed as mean ± S.E. of mean (SEM) of several replicates. The parallel control and formulation fed samples were compared by Holm Sidak posthoc test.

## 2 Results

### 2.1 AR feeding improved median survival of wild type flies maintained under crowded conditions

In order to examine effect of crowding stress on median life span, 50 or 100 larvae and flies were maintained per vial and the proportion of flies that survived on different days were compared with those maintained under uncrowded condition of 25 larvae/flies per vial. In agreement with earlier observations (Dwivedi et al. 2012), under uncrowded condition, AR feeding increased the median life span of flies when compared with those reared on normal food (**Table 1**). Crowding stress inflicted by keeping 50 or 100 larvae/flies per vial reduced the median life span of flies reared on regular as well as AR supplemented food (**Table 1**). Interestingly, however, the median life span of AR fed flies continued to be significantly greater than of those reared on normal food (**Table 1**). Further, while maximum crowding (100 flies/food vial) on normal food reduced the median life span by about 20% of the uncrowded condition, for the AR-fed flies the reduction was only about 9% of the uncrowded condition.

**Table 1.** Flies reared on AR supplemented food show better tolerance to crowding stress

| No. of flies per vial | Mean (±S.E.) Median life span of flies (in days) reared on different foods | |
| --- | --- | --- |
|  | Control | AR supplemented |
| **25** | 35.4 ± 1.1 | 41.2 ± 1.5* |
| **50** | 33.6 ± 1.8 | 39.8 ± 1.4* |
| **100** | 28.2 ± 1.3 | 37.6 ± 1.9* |

\* Values significantly different ( $P < 0.001$ ) from the corresponding control values N= 8 replicates of 25, 50 or 100 flies as per requirement.

### 2.2 AR feeding improved thermo-tolerance in young as well as older flies

It was reported earlier (Dwivedi et al. 2012) that compared to those fed on normal food, the AR fed third instar larvae and 3 day old flies displayed better tolerance to heat shock at 38^0^C for 60 minutes. In the present study, we examined if older flies also show better thermotolerance when reared on AR supplemented food. Thermotolerance of 3, 15, 30, 45 and 60 day old flies, reared since 1^st^ instar larval stage on AR supplemented or normal (control) food was assessed by survivorship after 24 h of the thermal stress and by comparing the proportion of flies knocked down after 15, 30, 45 or 60 min exposure to 38^0^C. As the data in Table 2 show, the proportion of flies surviving 24 h after the 60 min exposure to 38^0^C was significantly higher for the AR-fed flies than for those reared on normal food, although as expected, older flies showed greater sensitivity to 60 min exposure to 38^0^C. Similarly, compared to normal food reared flies, the proportion of flies that were knocked down at different time points of the heat shock was always lower for AR-fed flies, except for the 100% knock down observed in both sets of 45 and 60 day old flies thermally stressed for >30 min. These results thus show that even older flies have better thermo-tolerance when reared on AR supplemented food.

**Table 2.** Feeding on 0.5% AR supplemented food provides high degree of thermo-tolerance to the flies of different age groups

| Age | Duration (min)<br>at 38<sup>0</sup>C | Mean ± S.E. % flies<br>Knocked down | | Mean ± S.E. % flies<br>surviving after 24 h | |
| --- | --- | --- | --- | --- | --- |
|  |  | Control | AR | Control | AR |
| **3 Day** | **15** | 0 | 0 | 64.5 ± 4.2 | 79.5 ± 2.6* |
| | **30** | 2 ± 1.1 | 1.5 ± 1.0 | | |
| | **45** | 61 ± 2.6 | 32 ± 1.8* | | |
| | **60** | 83 ± 2.5 | 60 ± 1.4* | | |
| **15 Day** | **15** | 1.5 ± 0.7 | 1.5 ± 0.5 | 61.5 ± 3.6 | 78.5 ± 2.3* |
| | **30** | 3.5 ± 1.4 | 2 ± 1.1 | | |
| | **45** | 65 ± 2.5 | 28.5 ± 2.3* | | |
| | **60** | 84.5 ± 2.6 | 67 ± 2.1* | | |
| **30 Day** | **15** | 50.5 ± 2.4 | 29.5 ± 2.9* | 41.5 ± 2.3 | 70.5 ± 3.4* |
| | **30** | 83.5 ± 3.1 | 55.5 ± 2.4* | | |
| | **45** | 98.5 ± 1.1 | 81.5 ± 2.1 * | | |
|  | **60** | 100 | 100 |  |  |
| **45 Day** | **15** | 93.5 ± 2.0 | 70.5 ± 2.1* | 26 ± 2.1 | 48.5 ± 2.4* |
| | **30** | 100 | 91.5 ± 1.8* | | |
|  | **45** | 100 | 100 |  |  |
|  | **60** | 100 | 100 |  |  |
| **60 Day** | **15** | 100 | 92.5 ± 1.5* | 17 ± 1.9 | 36.5 ± 2.8* |
|  | **30** | 100 | 100 |  |  |
|  | **45** | 100 | 100 |  |  |
|  | **60** | 100 | 100 |  |  |

### 2.3 AR feeding reduced accumulation of Reactive Oxygen Species (ROS) in MT of 35 day old flies

It has been reported that ROS level increases as the flies age so that mutants deficient in ROS metabolism display reduced longevity and acute sensitivity to stress (Peng et al. 2014; Weber et al. 2012). Since, AR feeding resulted in increased median life span of wild type flies (Dwivedi et al. 2012), the ROS levels in larvae, young and aged flies reared on normal or AR supplemented food were estimated using the DCF fluorescence as indicator of ROS levels

MT from third instar larvae or 1 day old wild type flies fed on AR (**Figure 1A–F**) supplemented food did not display any significant difference in DCF fluorescence compared to their corresponding controls. Irrespective of the feeding regimen, the DCF fluorescence in larval and 1 day old adult MT was nearly absent or very low, respectively (**Figure 1A–F**). However, MT from 35 day old flies reared on normal food displayed very high DCF fluorescence (**Figure 1G, I**), indicative of accumulation of high levels of ROS as the flies age. Interestingly, MT from 35 day old flies reared on AR supplemented food since larval life showed significantly less DCF fluorescence (**Figure 1H, I**) indicating a reduced accumulation of ROS following dietary AR supplement. Examination of the DCF fluorescence in other tissues of larvae/flies also provided comparable results (not shown).

**Figure 1.**
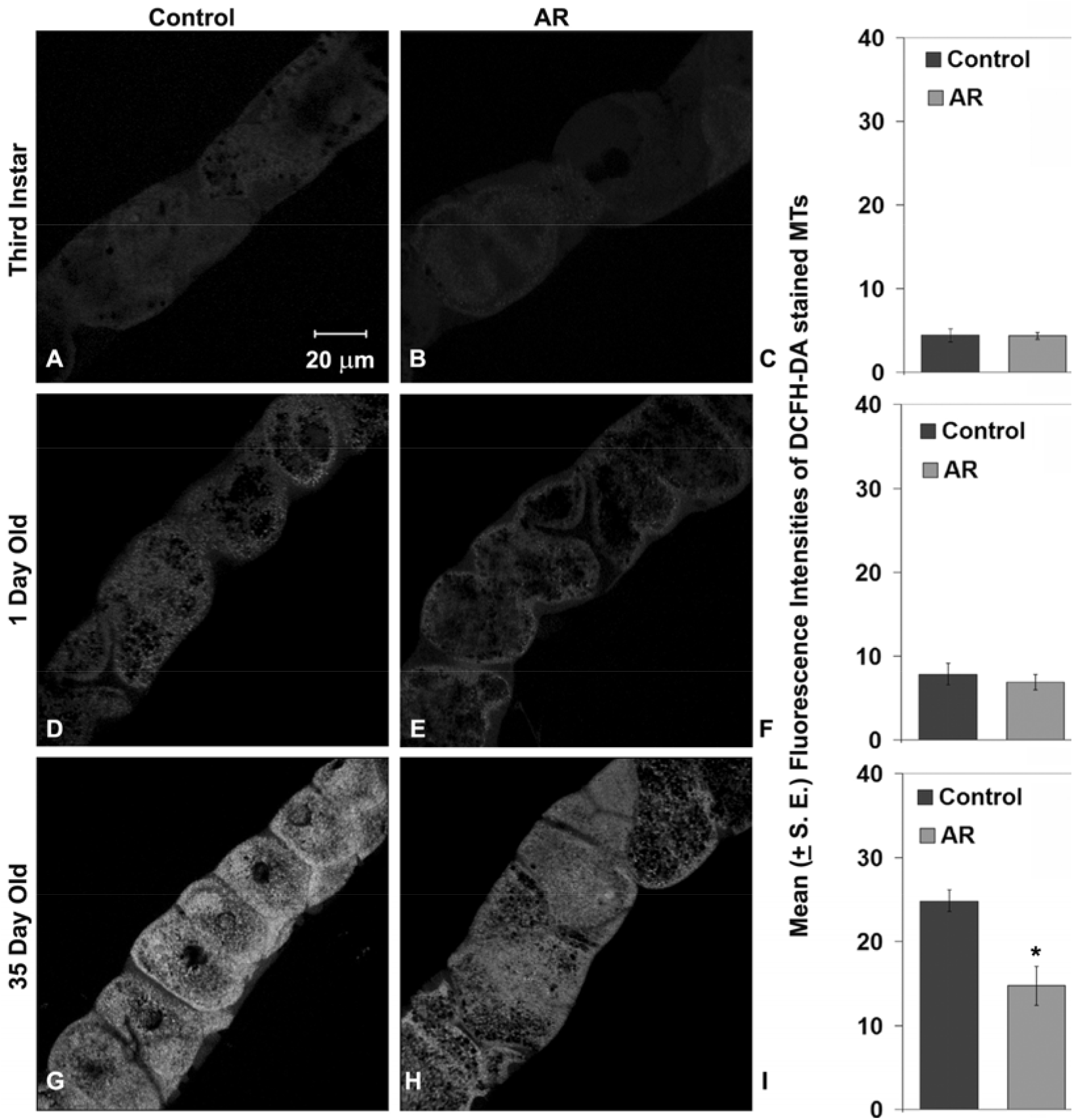
AR feeding reduced accumulation of ROS in MT cells in 35 day old flies. Confocal projections of four medial optical sections from MT of third instar larvae (**A, B**), 1 day old flies (**D, E**) or 35 day old flies (**G, H**) reared on normal (**A**, **D** and **G**) or AR (**B**, **E** and **H**) food, show DCF fluorescence following incubation in DCFH-DA. Scale bar in **A** represents 20µm and applies to all panels. It may be noted that as reported earlier (Dwivedi et al. 2012), MT in AR-fed individuals are wider. Histograms in **C, F** and **I** show mean (±S.E.) intensities of DCF fluorescence (N = 25 in each case); * in **I** indicates P < 0.01 when compared with the parallel control reared on regular food.

### 2.4 AR feeding provided better resistance to reactive oxygen species generated by paraquat in wild type flies

The N, N′-dimethyl −4, 4′-bipyridinium dichloride or paraquat is commonly used to generate reactive oxygen species (ROS) in vivo. Three day old wild type male flies, fed either on regular or AR supplemented food through their larval stages, were starved for six hours at 24^0^C and then transferred to vials without food but with a filter paper soaked in 10mM paraquat in 5% sucrose solution. The ROS accumulation in the flies following paraquat feeding was examined after 24 h using the DCF fluorescence (Cathcart et al. 1983) in Malpighian tubule (MT) cells. Compared to the intensity of DCF fluorescence in Malpighian tubules from paraquat treated flies reared on normal food, the signal was much less in tissues from similarly treated flies that were reared on AR supplemented food (**Figure 2A–C**).

**Figure 2.**
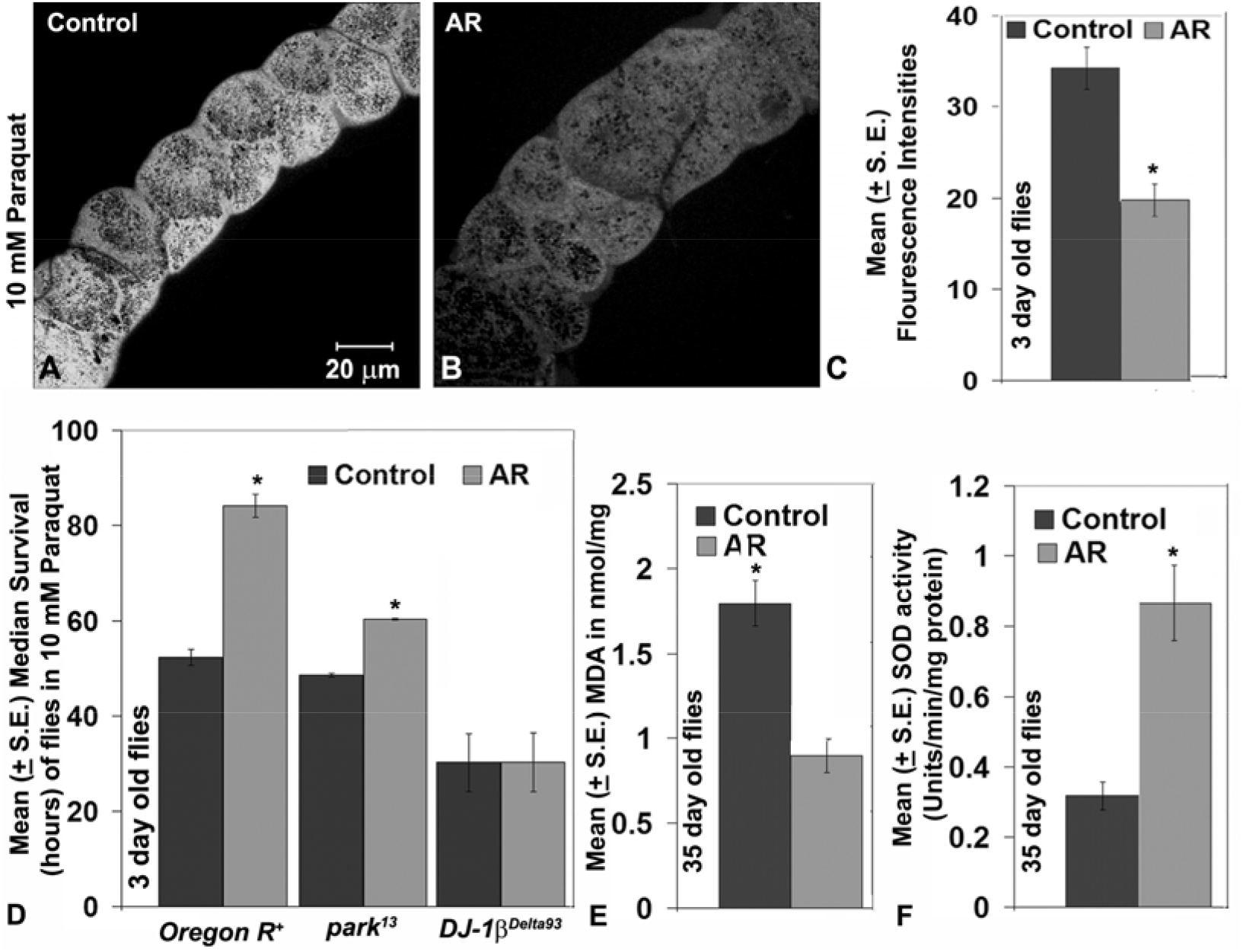
AR feeding improved the oxidative stress tolerance in wild type flies. **A** and **B** are confocal projections of four medial optical sections of MT showing DCF fluorescence in 3 day old flies treated for 24 h with 10mM paraquat followed by incubation in DCFH-DA. Scale bar in **A** represents 20µm and applies to images in **A** and **B**. Histogram in **C** shows mean (±S.E., N = 8 for each sample) intensities of DCF fluorescence in MT principle cells. Histogram in **D** shows mean (±S.E.) median survival of WT (*Oregon R*^+^), *park*^*13*^ and *DJ-1β*^*Delta93*^ flies (in hours, Y-axis) kept on 10mM paraquat (N for WT Control = 14 replicates, for AR = 12 replicates and for *park*^*13*^ and *DJ-1β*^*Delta93*^ Control and AR = 8 replicates of 25 male flies each). The bars in histogram shown in **E** represent mean (±S.E.) MDA levels (Y-axis) and those in **F** represent mean (±S.E.) SOD activity (Y-axis) in 35 days old flies fed either on regular food (Control) or AR supplemented (AR) food. * in **C**-**F** indicate the Means to be significantly different (P < 0.001) between corresponding Control and AR fed samples.

Effect of 10mM paraquat was also examined on survival of flies fed on regular or formulation supplemented food. In agreement with the reduced levels of ROS following paraquat exposure, the AR fed flies exhibited a significantly longer median survival (84.1 h) compared to the 52.4 h median survival of flies grown on regular food (**Figure 2D**). As a negative control, flies reared on normal or AR food showed little death (data not presented) when exposed in parallel to 5% sucrose solution.

### 2.5 AR feeding partially improved survival of *park*^*13*^ mutant flies but not of the *DJ-1β*-null flies exposed to paraquat oxidative stress

Mitochondrial ROS generation is implicated in neurodegenerative diseases like Parkinson’s disease (PD, Bonifati et al. 2003; Faust et al. 2009; Joselin et al. 2012; Munoz-Soriano and Paricio, 2011; Narendra et al. 2008, Yang et al. 2005; Yokota et al. 2003). ROS play a key role in the movement of Parkin into mitochondria in neurons (Joselin et al. 2012). A dominant null mutant allele of *parkin* in *Drosophila, park*^*13*^ (Greene et al. 2005), has been widely used as a fly model of PD. Another PD associated gene, *DJ-1 β*, has a role in protection from oxidative stress (Bonifati et al. 2003; Faust et al. 2009; Meulener et al. 2005; Yang et al. 2005; Yokota et al. 2003). The *park*^*13*^ as well as *DJ-1β*^*Delta93*^ individuals are reported to be highly sensitive to oxidative stress (Botella et al. 2009; Meulener et al. 2005). Therefore, *park*^*13*^/*TM6B* and *DJ-1β*^*Delta93*^ larvae were reared on regular or AR supplemented food and the 3 day old male flies were exposed to 10mM paraquat treatment and monitored every 12 hour to count the number of surviving flies. In agreement with earlier studies (Joselin et al. 2012), the *park*^*13*^/*TM6B* flies reared on regular food were sensitive to paraquat. AR feeding significantly improved the survival of paraquat treated *park*^*13*^/*TM6B* flies although the improved median survival was less than that of the AR reared wild type flies exposed to paraquat (**Figure 2D**). Interestingly, however, AR feeding did not improve the survival of *DJ-1β*^*Delta93*^ (*DJ-1β* null) flies following exposure to paraquat (**Figure 2D**). Formulation fed or regularly fed flies kept in parallel on 5% sucrose solution did not show any significant death during the period of observation (not shown). These results indicate that *DJ-1β*, is essential for AR mediated rescue from paraquat-induced oxidative stress.

### 2.6 AR feeding decreased lipid peroxidation and increased SOD activity in flies

Aging is associated with an increase in oxidative damage to nucleic acids, sugars, sterols and lipids (Lushchak, 2014; Peng et al. 2014; Rikans and Hornbrook, 1997; Sies, 2015; Zhao and Haddad, 2011). Since lipid peroxidation is one of the markers for age-associated oxidative damage (Kasapoglu and Ozben, 2001, Peng et al. 2014), we estimated levels of lipid peroxidation in formulation fed and regularly fed 35 day old flies. Lipid peroxide levels were expressed as nano-moles of malondialdehyde (MDA) formed per mg of protein following exposure of the fly homogenates to the assay mixture containing sodium dodecyl sulphate and thiobarbituric acid (Ohkawa et al. 1979). Compared to the regularly fed flies, AR fed 35 day old flies showed significantly less concentration of MDA (**Figure 2E**) reflecting reduced lipid peroxidation.

An inadequate clearance of ROS like superoxides, hydrogen peroxide and the hydroxyl radicals results in oxidative damage, leading to physiological dysfunction and many types of age related disorders/diseases (Chaudhari et al. 2007; Park et al. 2005; Tsuzuki et al. 2007; Turner at al. 2009). Glutathione-dependant antioxidants and enzymes like superoxide dismutase (SOD), catalase (CAT) etc. have an evolutionarily conserved role in the metabolism and removal of ROS (Muller at al. 2007). We compared the levels of Cu–Zn SOD activity in 35 day old wild type flies reared on AR supplemented or normal food. Interestingly, AR feeding remarkably enhanced the Cu–Zn SOD activity (**Figure 2F**) in aged flies when compared to those reared in parallel on normal food (**Figure 2F**).

### 2.7 AR feeding did not affect the Hsp70/Hsc70 and Hsp83 expression under normal conditions, on heat shock or during recovery but enhanced Hsp27

Since different stress-induced heat shock proteins have been implicated in conferring stress-tolerance and modulating life span (Arya et al. 2007; Feder and Hoffmann, 1999; Lindquist, 1986; Landis et al. 2012; King and MacRae, 2015), we examined levels of Hsp70, Hsp83 and Hsp27 in unstressed and heat shocked tissues of larvae to see if their levels are affected by rearing on AR supplemented food.

Hsp70 is the most strongly induced heat shock protein in *Drosophila* following cell stress (Arya et al. 2007; Feder and Hoffmann, 1999; Guertin et al. 2010; Peterson and Lindquist, 1988). Therefore, expression of Hsp70 was examined in wild type third instar larval tissues reared on AR or regular food when maintained at 24^0^C or subjected to heat shock at 37^0^C for 1h or allowed to recover from the heat shock for 2 h. Levels of hsp70 transcripts in tissues from AR fed larvae, determined by real time PCR, did not show any significant difference from the corresponding samples of the regular food reared larvae (**Figure 3A**). Third instar larval salivary glands (SG) were immunostained with the stress-induced Hsp70 specific 7Fb antibody (Velazquez and Lindquist, 1984) following heat shock at 37^0^C or after 2h recovery from heat shock. SG from AR or regular food fed larvae showed comparable distribution of Hsp70 (**Figure 3B, B’, C, C’**). Unstressed cells from regular or AR fed larvae did not show any detectable levels of Hsp70 (not shown). Western blot detection of Hsp70, using the 7Fb antibody, in whole larval proteins from wild type larvae fed on regular or AR supplemented food and subjected to heat shock at 37^0^C or allowed to recover at 24^0^C from the 1 h heat shock confirmed the above immunostaining results that AR feeding did not alter the levels of induced Hsp70 protein on heat shock or during recovery (inset in **Figure 3B’, C’**).

**Figure 3.**
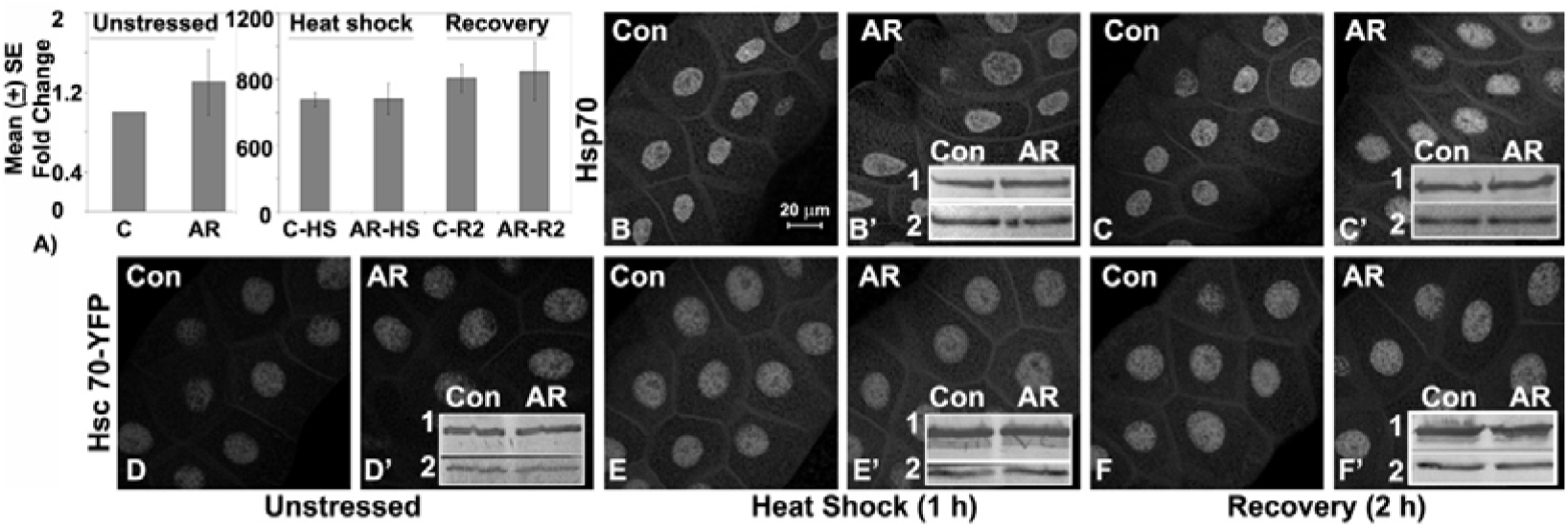
Feeding larvae on AR supplemented food did not affect the distribution and levels of Hsp70/Hsc70 on heat shock and during recovery. **A**- Histograms show mean (± S.E.) fold changes of Hsp70 transcripts (Y-axis) in larvae fed on normal (**C**) or AR (**AR**) food at 24^0^C (unstressed) or following heat shock (**C/AR-HS**) or after 2 h recovery from heat shock (**C/AR-R2**); the Hsp70 mRNA levels in unstressed larvae reared on normal food (**C**) were taken as 1 for estimating the fold changes in other samples. **B-F’**-Confocal projection images showing levels of Hsp70 (**B-C’**) or Hsc70Cb-YFP (**D-F’**) in salivary glands from third instar larvae reared on control (**Con**) or AR supplemented (**AR**) food under different conditions as noted at the bottom (Hsp70 immunostaining for unstressed condition is not shown). The scale bar in **B** (20 μm) applies to **B**-**F’**. Insets in **B’, C’** are western blots showing levels of Hsp70 detected by the 7Fb antibody (rows **1**) and those in **D’-F’** show levels of Hsc70/Hsp70 detected by the 7.10.3 antibody (rows **1**) in total proteins from differently fed (**Con** and **AR**) wild type late 3^rd^ instar larvae under unstressed (**D’**) or after heat shock (**B’**, **D’**) or after 2 h recovery from heat shock (**C’**, **F’**); β-tubulin levels detected by the E7 antibody, used as the loading control, are shown in lower rows (**2**) in each western blot in the insets.

We also examined if the expression of YFP-tagged Hsc70Cb, one of the constitutively expressed Hsc70 family proteins, is affected by AR feeding. The pattern and intensity of YFP fluorescence in SG from late *Hsc70Cb-YFP* larvae fed on regular or AR food did not differ from each other neither under normal conditions nor after heat shock or during recovery from heat shock (**Figure 3D, D’, E, E’, F, F’**). Western blot detection of all Hsc70/Hsp70, using the 7.10.3 Ab, which detects all constitutive and inducible members of the Hsp70 family in *Drosophila* (Palter et al. 1986), in total proteins from wild type larvae reared either on AR or on regular food confirmed that the levels of Hsc70/Hsp70 remained similar under all conditions (insets in **Figure 3D’, E’, F’**).

In unstressed cells Hsp83 was abundant in cytoplasm but much less in nuclei of SG from larvae reared on normal or AR supplemented food (**Figure 4A**, **D**). Following heat shock (**Figure 4B**, **E**) as well as during recovery from heat shock (**Figure 4C**, **F**), the Hsp83 was more abundant in SG nuclei than in cytoplasm. However, under all the three conditions, the distribution pattern and levels of Hsp83 in SG polytene cells from AR supplemented food reared larvae were not different from those in normal food reared larvae (**Figure 4**). This was also confirmed by quantification of Hsp83 fluorescence intensity in different samples (**Figure 4G**). Similar results were obtained with MT and eye disc cells (not shown). Levels of hsp83 transcripts in differently fed larvae were also examined under unstressed conditions, after heat shock and after 2 h recovery by real time PCR. It was observed that levels of *hsp83* transcripts in larvae reared on normal food and AR supplemented food were comparable under various conditions (**Figure 4H**).

**Figure 4.**
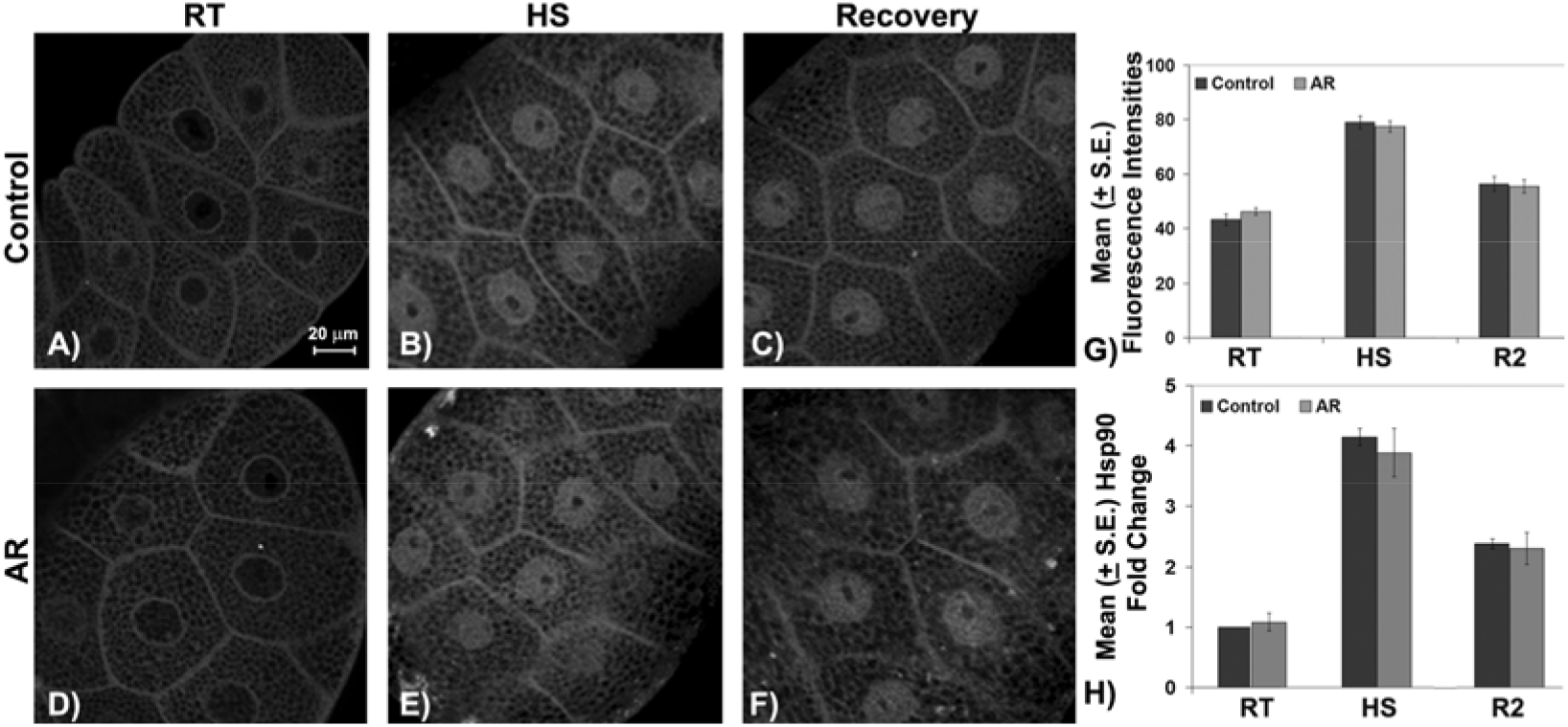
Feeding larvae on AR supplemented food did not affect the distribution and levels of Hsp83 protein and transcripts. **A**-**F** are confocal projections of four medial optical sections of third instar larval salivary glands immunostained with anti-Hsp83 antibody either at 24^0^C (unstressed, **A, D**), or after heat shock at 37^0^C for 1 hour (**B, E**) or after 2 h recovery from heat shock (**C, F**). Scale bar in **A** represents 20µm and applies to images **A-F**. Histogram in **G** shows the mean (± S.E.) fluorescence intensities of Hsp83 in SG of third instar larvae reared on normal (Control) or AR supplemented (AR) food and subjected to various conditions. **H**- Histogram represents mean (± S.E.) fold change of Hsp83 transcripts in larvae fed on AR or normal food either kept under normal condition or heat shocked (HS) or recovered for 2 hours (R2).

The Hsp27 protein is normally distributed in nuclei as well as in cytoplasm of wild type SG from larvae reared on regular food (**Figure 5A**). Following heat shock, Hsp27 levels are enhanced in both nucleus as well as cytoplasm (**Figure 5B**), which nearly restored to that in unstressed cells after 2 h recovery from heat shock (**Figure 5C**). Interestingly, SG cells from AR fed larvae showed significantly higher levels of Hsp27 in nucleus as well as cytoplasm of unstressed cells than in those from regular food reared larvae (compare **Figure 5A** and **D**). Heat shock further elevated levels of Hsp27 in AR fed larval SG (**Figure 5E**). Unlike in regular fed larval SG cells, Hsp27 levels in AR larval SG remained higher even after 2 hour recovery from heat shock (**Figure 5C** and **F**). The above noted differences in levels of Hsp27 between normal and AR supplemented food confirmed by quantification of the fluorescence intensity of immunostained cells (**Figure 5G**). Other tissues like MT, eye and other imaginal discs also displayed a comparable difference in the levels of Hsp27 in regularly fed or AR fed larvae (not shown). Examination of levels of hsp27 transcripts by real time PCR showed that in parallel with the immunostaining patterns, hsp27 transcripts were significantly more abundant under all conditions (unstressed, heat shock and 2 h recovery) in AR fed larvae than in those reared on regular food. Heat shock resulted in greater induction of hsp27 transcripts in AR while after 2 hour recovery, the hsp27 transcripts in AR fed samples still remained higher than that in normal fed recovery sample (**Figure 5H**).

**Figure 5.**
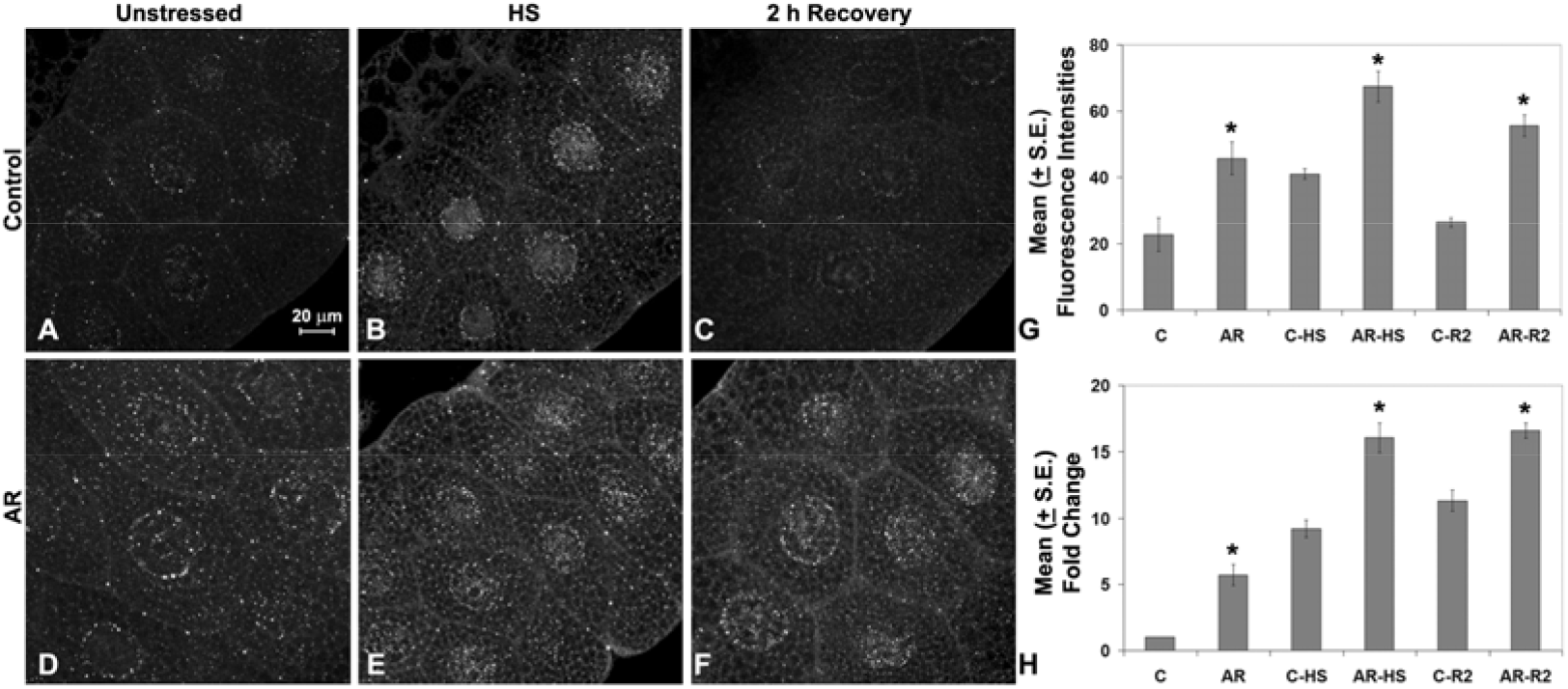
Feeding larvae on AR supplemented food elevated the levels of Hsp27 protein and transcripts. **A**-**F** are confocal projections of four medial optical sections of third instar larval salivary glands immunostained with anti-Hsp27 maintained at 24^0^C (unstressed, **A, D**), or after heat shock at 37^0^C for 1 h (**B, E**) or after 2 h recovery from heat shock (**C, F**). Scale bar in **A** represents 20µm and applies to images **A-F**. Histogram in **G** shows the mean (± S.E.) fluorescence intensities of Hsp27 (Y-axis) in SG of third instar larvae reared on regular (C) or AR supplemented (AR) food subjected to various conditions (X-axis). **H**- Histogram represents mean (± S.E.) fold change in levels of Hsp27 transcripts in larvae fed on regular (C) or AR supplemented (AR) food (X-axis). In **G** and **H**, **C** and **AR** indicate unstressed larvae reared on regular or AR supplemented food, **HS** indicate heat shocked samples while **R2** indicates samples recovered for 2 h from the heat shock. * indicates P < 0.001 when compared with the corresponding control (**AR** vs. **C**).

## 3 Discussion

Ayurvedic system, the oldest among various traditional health care practices in world, is claimed to provide a holistic health care (Patwardhan et al, 2015; Singh, 2009; Lakhotia, 2016). Ayurveda has its own wide range of formulations consisting of herbal, mineral and other biological materials which have been compiled as ‘*Dravyaguna Sangrah’* (Sharma, 1994; Singh, 2009). However, we know very little about the specific modes of actions of the various complex Ayurvedic formulations (Valiathan, 2006, Lakhotia, 2013, 2016). Natural products present in Ayurvedic formulations have provided interesting sources of new molecular entities with potential therapeutic applications. This has stimulated a large number of studies on various herbs to identify the ‘active principles’. However, Ayurvedic herbal and other *Rasayana*s are complex mixtures of multiple ingredients which work in a synergistic manner and, therefore, the reductionist approach to identify the ‘active principle’ does not always provide information on the mechanisms through which the classical formulations exert their effects (Lakhotia, 2013, 2016; Patwardhan et al, 2015).

Earlier studies from our laboratory (Dwivedi et al. 2012, 2013, 2015) established *Drosophila* as a model organism to understand the mechanisms and modes of actions of the complex Ayurvedic formulations. As noted earlier, one of the major usages of AR in humans is to rejuvenate and thus improve longevity and maintain youthfulness. Earlier studies showed that feeding flies on 0.5% AR supplemented food enhanced their life span, improved fecundity and enhanced cellular levels of key regulatory molecules like the hnRNPs, CBP300 (Dwivedi et al. 2012), suppressed neurodegeneration associated with polyQ disorders and Alzheimer’s Disease (Dwivedi et al. 2013), and prevented induced apoptosis without affecting developmental cell death (Dwivedi et al. 2015). A study in mouse model has shown that the genomic integrity in aging mice provided with AR is significantly better than in control sibs (Swain et al. 2012). Present studies provide further mechanistic insights into the reported beneficial and rejuvenation affects of dietary AR.

Besides the earlier reported improved tolerance of AR-fed flies to starvation and thermal stress (Dwivedi et al. 2012), present studies clearly show that AR feeding significantly improves tolerance to crowding and oxidative stresses. A more significant finding is that AR-feeding makes even older wild type flies more tolerant to a severe heat shock (38^0^C). Such improved capability to face various external environmental stresses as well as internal metabolic oxidative stresses seem to promote the longer life span displayed by the AR fed flies. The improved starvation-tolerance (Dwivedi et al. 2012) and resistance to crowding displayed by AR-fed larvae/flies and their increased longevity agrees with the earlier reported correlation that better larval physiology promotes better starvation tolerance and longevity of adults (Chippindale et al. 1996; Mueller and Barter, 2015; Prasad and Joshi, 2003; Service 1987).

Several studies have reported very strong anti-oxidant activity of extracts of Amla or Indian Gooseberry fruit, the principal component of AR (Bargale et al. 2014; Chatterjee et al. 2011; Govindarajan et al. 2004; Khan, 2009; Krishnaveni and Mirunalini, 2010; Poltanov et al. 2009; Samarakoon et al, 2011; Scartezzini and Speroni, 2000). In agreement, present in vivo results show that AR feeding significantly improved the oxidative stress tolerance of wild type flies and remarkably reduced the accumulation of ROS with age, reflecting an improved cellular redox homeostasis. The partially improved survival of *park*^*13*^ flies but no improvement in survival of paraquat exposed *DJ-1β* mutant flies shows that the improved management of ROS by AR feeding requires functional Parkin but more so the DJ-1β protein. The DJ-1β protein plays a significant role in managing oxidative stress and the Phosphatidylinositol 3-Kinase/Akt signalling pathway (Meulener et al. 2005; Yang et al. 2005).

Aging is strongly correlated with sensitivity to components that increase the oxidative stress and thus cause greater damage to macromolecules like DNA and lipids (Haigis and Yanker, 2015; Holmbeck et al. 2014; Lagouge et al. 2013; Mallikarjun et al. 2014; Weber et al. 2012). AR fed flies had reduced levels of ROS and reduced levels of lipid peroxidation. Populations of flies with longer life span exhibit higher frequencies of the high-activity allele of the Cu–Zn SOD which has an important role in conferring resistance to oxidative stressors through removal of superoxides (Peng et al. 2014; Tyler et al. 1993). The significantly improved SOD activity in AR fed older flies contributes to reduced oxidative damage, which in turn would improve fecundity and life span. The increased resistance to oxidative stress following AR feeding appears to also underlie the reported reduced DNA damage seen in aged mice (Swain et al, 2012).

The significantly greater thermotolerance exhibited by older flies that were reared on AR supplemented food is remarkable. Our results show that unlike the generally strong correlation (Krebs and Feder, 1998; Lin et al. 2014; Stetina et al. 2014) between thermotolerance and expression of heat shock proteins, especially the Hsp70, AR did not significantly affect expression of Hsp83 and Hsp70 proteins or transcripts. On the other hand we found that basal as well as heat shock induced levels of Hsp27 were significantly more elevated in AR fed larvae. It is interesting that although Hsp27 is reported to be unessential for surviving heat shock (Hao et al. 2007), it plays significant roles in tolerance to other insults like starvation, oxidative stress etc and also in determining the life span of flies (Hao et al. 2007, Pandey et al. 2015; Shi et al. 2011; Wang et al. 2004). Thus, the improved thermo-tolerance of AR fed larvae and flies does not appear to be due to their stronger heat shock response. It may be noted that the heat shock response is a transient process and its long-term activation may actually be harmful (Lamech and Haybes, 2014). The improved thermo-tolerance of AR fed flies may thus be due to better management of oxidative stress and the overall improved physiology and homeostasis (Dwivedi at al., 2012, 2013, 2015). AR supplement may promote these beneficial changes through enhanced levels of key regulatory molecules like different hnRNPs and CBP300 (histone acetyl transferase) and proteasomal activity that have global roles in gene expression. Various environmental insults (external as well as internal) also induce cells to undergo apoptotic, which is one of the factors underlying aging (Lu et al. 2012; Pérez-Garijo and Steller, 2015). Our earlier finding that AR feeding suppresses induced but not the developmental apoptosis also indicates a better cellular tolerance promoted by AR to environmental insults (Dwivedi et al. 2015).

## 4 Conclusions

The modern medicine and Ayurveda, although have universal attributes and share the common objective of well being of mankind, seem to differ in their philosophical and epistemological foundations, conceptual framework and practical outlook (Patwardhan et al. 2015; Lakhotia, 2016). Most contemporary scientific studies to understand specific Ayurvedic formulations/therapies employ the typical reductionist approach of molecular biology to identify the so called “active principle”. However, this is not in consonance with the holistic and combinatorial approach of Ayurveda (Lakhotia, 2013, 2016; Patwardhan et al. 2015). Therefore, unbiased and a rational understanding of Ayurvedic Biology in a holistic manner is expected to provide better quality control of the various formulations.

*Drosophila* and vertebrates share a large number of genes, possess similar basic metabolic functions that maintain sugar, lipid and amino acid homeostasis (Baker and Thummel, 2007; Britton et al. 2002; Engelman, 2006; Leopold and Perrimon, 2007; Saltiel and Kahn 2001); they also share the highly conserved PI3K/Akt/mTOR signalling pathways that play central roles in regulation of oxidative metabolism and aging in both groups (Wang et al. 2012). In view of these and in view of the unparalleled genetic tractability, the fly model is increasingly used for understanding human health issues (Perrimon et al. 2016). Therefore, our findings in the fly model have clinical implications since the more robust physiological state following dietary AR supplement in flies suggest that AR may prevent several diseases and aging conditions and thus contribute to “healthy aging” in mankind.

## 5 Acknowledgements

We thank Arya Vaidya Sala, Kottakal (Kerala, India) for providing the *Amalaki Rasayana* formulation and Prof. M. S. Valiathan for initiating the coordinated studies on Science of Ayurveda. We thank Dr. P. K. Tiwari (Gwalior, India) for providing the anti-Hsp27, Dr. R. M. Tanguay (Canada) for anti-Hsp90 antibody and Dr. M. B. Evgen’ev (Russia) for 7Fb and 7.10.3 antibodies. Bloomington stock centre is acknowledged for providing various stocks used in present study. This work was supported in part by a grant (no. Prn.SA/ADV/Ayurveda/6/2006) from the Office of the Principal Scientific Advisor to Govt. of India (New Delhi) and by the Raja Ramanna Fellowship of the Department of Atomic Energy (Govt. of India, Mumbai) to SCL. The Confocal Facility, established by the Department of Science & Technology, Govt. of India (New Delhi), is supported by the Banaras Hindu University. VD is supported by research fellowship from University Grants Commission (New Delhi).

